# How individual *P. aeruginosa* cells with diverse stator distributions collectively form a heterogeneous macroscopic swarming population

**DOI:** 10.1101/2023.04.10.536285

**Authors:** Jaime de Anda, Sherry L. Kuchma, Shanice S. Webster, Arman Boromand, Kimberley A. Lewis, Calvin K. Lee, Maria Contreras, Victor F. Medeiros Pereira, Deborah A. Hogan, Corey S. O’Hern, George A. O’Toole, Gerard C.L. Wong

**Affiliations:** Department of Bioengineering, University of California Los Angeles, CA 90095; Department of Chemistry and Biochemistry, University of California Los Angeles, CA 90095; California NanoSystems Institute, University of California Los Angeles, CA 90095; Department of Microbiology and Immunology, Geisel School of Medicine at Dartmouth, Hanover, New Hampshire, United States of America; Department of Mechanical Engineering & Materials Science, Yale University, New Haven, CT 06520 USA; Department of Nanoengineering, University of California San Diego, CA 92093

## Abstract

Swarming is a macroscopic phenomenon in which surface bacteria organize into a motile population. The flagellar motor that drives swarming in *Pseudomonas aeruginosa* is powered by stators MotAB and MotCD. Deletion of the MotCD stator eliminates swarming, whereas deletion of the MotAB stator enhances swarming. Interestingly, we measured a strongly asymmetric stator availability in the WT strain, with MotAB stators produced ∼40-fold more than MotCD stators. However, recruitment of MotCD stators in free swimming cells requires higher liquid viscosities, while MotAB stators are readily recruited at low viscosities. Importantly, we find that cells with MotCD stators are ∼10x more likely to have an active motor compared to cells without, so wild-type, WT, populations are intrinsically heterogeneous and not reducible to MotAB-dominant or MotCD-dominant behavior. The spectrum of motility intermittency can either cooperatively shut down or promote flagellum motility in WT populations. In *P. aeruginosa*, transition from a static solid-like biofilm to a dynamic liquid-like swarm is not achieved at a single critical value of flagellum torque or stator fraction but is collectively controlled by diverse combinations of flagellum activities and motor intermittencies via dynamic stator recruitment. Experimental and computational results indicate that the initiation or arrest of flagellum-driven swarming motility does not occur from individual fitness or motility performance but rather related to concepts from the ‘jamming transition’ in active granular matter.

**Importance:** After extensive study, it is now known that there exist multifactorial influences on swarming motility in *P. aeruginosa*, but it is not clear precisely why stator selection in the flagellum motor is so important or how this process is collectively initiated or arrested. Here, we show that for *P. aeruginosa* PA14, MotAB stators are produced ∼40-fold more than MotCD stators, but recruitment of MotCD over MotAB stators requires higher liquid viscosities. Moreover, we find the unanticipated result that the two motor configurations have significantly different motor intermittencies, the fraction of flagellum-active cells in a population on average, with MotCD active ∼10x more often than MotAB. What emerges from this complex landscape of stator recruitment and resultant motor output is an intrinsically heterogeneous population of motile cells. We show how consequences of stator recruitment led to swarming motility, and how they potentially relate to surface sensing circuitry.

## Introduction

At a macroscopic level, populations of bacteria can abruptly organize into a motile ‘swarm’ (1, 2), but it is not clear how this process is collectively initiated or arrested. The underlying molecular mechanisms that underpin swarming motility are complex and heterogeneous. In swarming as in swimming, the flagellar motor provides propulsion (1). The basic flagellar structure is well known, with four main components: the extracellular helical filament, the hook, the rotor, and the stators. When the flagellum is active, the stators transform an ion flux across the cytoplasmic membrane into torque to rotate the rotor. Bacterial species have evolved diverse specialized flagellar motors with differences in 1) the number and type of stators (H^+^-vs Na^+^-powered) (3–5), 2) the diameter of the C-ring rotor (6), 3) presence or absence of a periplasmic ring (7), and 4) the number of flagella expressed (1, 8). Single stator systems such as that of the multi-flagellated enteric *Escherichia coli* recruit up to 11 H^+^-driven MotAB stators to power its flagella (9, 10), while the marine single-flagellated *Vibrio cholerae* uses up to 13 Na^+^-driven PomAB stators (6). Dual stator systems, such as those for *Shewanella oneidensis* MR-1 (4, 11) and multi-flagellated *Bacillus subtilis* (12), afford these microbes the versatility to utilize two ion gradients by interchangeably recruiting H^+^-driven or Na^+^-driven stators.

How stators are organized with respect to function in *Pseudomonas aeruginosa* is not as clear: *P. aeruginosa* employs a dual H^+^-driven stator system, MotAB and MotCD, but does not exploit different ion gradients, so it is unclear why this apparent stator redundancy exists. In fact, the two stators look remarkably similar in terms of amino acid sequence and do not have significantly different torque outputs per stator unit (5). Despite these similarities, the MotAB and MotCD stator sets have been shown to produce remarkably different motility phenotypes. It has been observed previously that a strain with a exclusively MotAB-powered motor can swim faster than its MotCD-powered motor counterpart strain (13, 14), whereas a strain with the MotCD-powered motor forms a significantly larger swarm area than a strain with only the MotAB stator (15, 16), suggesting these stator sets possess distinct functional capabilities despite their noted similarities. Indeed, it is also not clear how *P. aeruginosa*, a monotrichous species, can swarm so efficiently given that most other bacteria that exhibit swarming motility are polytrichous species, e.g. *E. coli, B. subtilis,* or *Salmonella enterica* (1).

Here we examine how *P. aeruginosa* uses different two-stator configurations to initiate or arrest flagellum-driven motility collectively in a population, and thereby control swarming behavior. The root phenomenon that enables control of collective flagellum driven motility and environmentally sensitive responses is a biased adaptive stator recruitment mechanism. To facilitate high swim speeds in low viscosity environments (i.e., swimming), the flagellar motor is primarily decorated with MotAB stators, while high viscosity environments (i.e., swarming) promote MotCD recruitment, but such recruitment is hindered by high levels of MotAB production. Our data suggest that the torque outputs of motors driven by MotAB or MotCD are not markedly different at high viscosity, consistent with other measurement modalities (5). We find the unanticipated result that the two motor configurations have significantly different motor intermittencies, that is, the fraction of flagellum-active cells in a population on average. What emerges from this complex landscape of stator recruitment and resultant motor output is an intrinsically heterogeneous population of motile cells not readily reducible to MotAB-dominant or MotCD-dominant behavior. Based on experimental and computational data, we find that the initiation of flagellum-driven swarming activity occurs when numerous individual motility choices, achieved via stator recruitment, are successfully integrated into macroscopic community motion via processes related to the ‘unjamming’ transition in the field of ‘granular matter’; which separates static, solid-like, sessile behavior from flowing, liquid-like, motile behavior in a population of cells. Conversely, the sudden and collective arrest of flagellum-driven motility can be achieved via changing the composition of stators in the flagellum motor, even for a relatively small subpopulation of cells in a heterogeneous population. The conceptual results presented here are in principle generalizable. Beyond an improved understanding of flagellum-driven swarming phenomena, a collective shutdown of flagellum motility orchestrated by stator recruitment may be important to surface sensing-mediated signaling pathways that lead to flagellum shutdown for heterogeneous bacterial populations during the initiation of biofilm formation.

## Results

### The WT and MotAB motor exhibit greater swimming speed than the MotCD motor in low viscosity environments, but all strains have similar swimming speed in high viscosity environments

A natural starting point of comparison between the output of WT as well as MotAB- and MotCD-exclusive flagellar motors is swimming speeds. Here it is illuminating to compare single cell swimming speeds in low viscosity fluids, but also high viscosity environments typically encountered in swarming. Note, that for this manuscript, when we refer to the role of the MotAB motor, we are measuring the behavior of a Δ*motCD* mutant, which lacks MotCD motor function. In contrast, when we refer to a MotCD motor, this shorthand indicates a Δ*motAB* mutant which lacks any MotAB motor function. The wild-type (WT) motor has both functional MotAB and MotCD stators (Fig. 1A).

**Figure 1.**
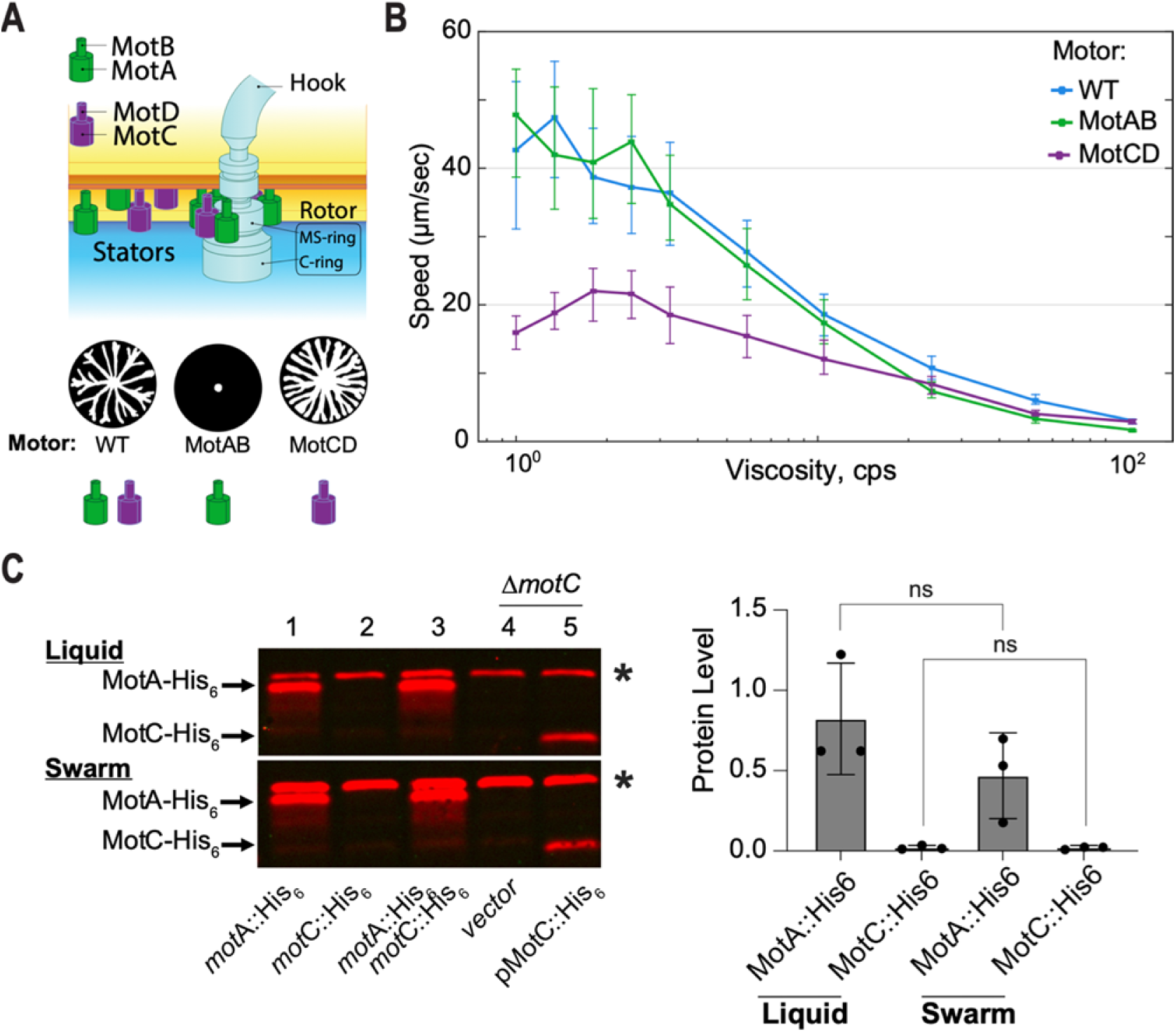
Convergence of swimming speeds by the MotCD-motor to the WT- and MotAB-motors with increasing viscosity and asymmetric production of stator-type sets in WT. *(A)* The dual proton-driven stator system, MotAB and MotCD, of *Pseudomonas aeruginosa.* Stators are recruited to the motor to provide the torque necessary to rotate the flagella. Three motor configurations can be formed: the swarming MotABCD (WT) motor, the swarming deficient MotAB motor, and hyper-swarmer MotCD motor. A representative swarming pattern for each motor type is presented above each stator set. *(B)* Measurement of swimming speed for three stator configurations: two-stator WT (MotABCD), and the MotAB and MotCD single-stator motors at different viscosities (by increasing the percent concentration of methylcellulose in solution). At least 100 trajectories per viscosity condition were measured. Error bars denote the first and third quartiles of the distribution about the mean. *(C)* Western blot detection of the MotA::His_6_ and MotC::His_6_ epitope-tagged proteins expressed in membrane fractions of the strains indicated. In lanes 1 through 3, the proteins are expressed from the respective endogenous loci under native promoter control. Samples in lane 3 derive from the strain in which both the *motA* and *motC* genes express the respective proteins fused to the His_6_ epitope. In lanes 4 and 5, samples from the Δ*motC* strain harbor either the empty vector control (lane 4) or a multi-copy plasmid for expression of the MotC::His_6_ protein under arabinose induction via the P_bad_ promoter. Arrows point to the location of the indicated proteins. The asterisk (*) indicates a non-specific band present in all samples and used as a normalization control for quantification. Strains were grown for 16h in either liquid (top panel) or swarm agar plates (bottom panel) with 0.05% arabinose to induce plasmid-borne MotC::His_6_ expression. Proteins were detected using an anti-His_6_ monoclonal antibody. Bar plot on the right show the quantification of the MotA::His_6_ and MotC::His_6_ proteins from three biological replicates of samples from the *motA*::His_6_ *motC*::His_6_ double-tagged strain grown under the indicated conditions (panel d, lane 3, top and bottom). Data were analysed by one-way ANOVA followed by Tukey’s post-test comparison. ns, not significantly different.

We find that the WT and the MotAB motor facilitated faster swimming than the MotCD motor at low viscosities. In aqueous medium, the linear speed of the MotAB motor (48.5 ± 16.2 μm/sec) outperforms that of the MotCD motor by 3-fold (15.0 ± 5.0 μm/sec). However, as viscosity is increased via addition of methylcellulose (17–19), the swim speeds decrease. Importantly, the swim speeds of the three strains converge as extracellular viscosity increases (Fig. 1B): At ∼24 cP, the difference in swim speed between the MotAB and MotCD motor is not significant (7.2 ± 3.5 and 8.1 ± 2.9 μm/sec, respectively). These data indicate that at low viscosity, the MotAB motor is capable of faster swimming than the MotCD motor, although at high viscosity, both motors perform similarly, a conclusion supported by a recent report (5).

### Production of MotAB and MotCD stators is strongly asymmetric, likely enforcing uneven competition for motor recruitment in WT

For the two-stator system of *P. aeruginosa,* the quantity as well as the type of stator is crucial to understanding motility. While the WT motor is generally thought to utilize MotCD stators to facilitate swarming (20, 21); given that, it is not clear how the MotCD motor strain significantly outperforms the WT and MotAB motor strains in swarming motility. Here, we test the hypothesis that the relative levels of MotAB and MotCD stator pairs in the cell modulate stator composition of the motor.

To answer this question, we measured levels of the MotA and MotC proteins within the same population of cells by inserting a His_6_ epitope tag into the C-termini of the stator proteins encoded by the *motA* and *motC* genes at their respective loci on the chromosome of an otherwise WT strain. This strain allows detection of both the MotA-His (31 kDa) and MotC-His (26.8 kDa) proteins expressed under native promoter control in the same cells via the same antibody, but distinguishable by their molecular weight difference. Given that the MotA and MotB subunits are co-expressed in an operon, as are the MotC and MotD subunits, the MotA::His_6_ and MotC::His_6_ protein levels serve as a proxy for the levels of their respective stator pairs, MotAB and MotCD.

As shown in Fig. 1C levels of the MotA::His_6_ protein (lanes 1 and 3, left panel) and MotC::His_6_ protein (lanes 2 and 3) are not significantly different between liquid and swarm growth conditions (Fig. 1C, right panel). Note that lane 3 samples represent the *motA*::His_6_ *motC*::His_6_ strain bearing both epitope tags. The MotA::His_6_ protein levels are strikingly higher than those of MotC::His_6_ (Fig. 1C; lane 3 and right panel) - MotC::His_6_ is visually difficult to detect, regardless of whether grown in liquid or on swarm plates. There is ∼40-fold more MotA::His_6_ protein compared to MotC::His_6_.

The low level of MotC is not likely due to destabilization by insertion of the His epitope tag, as we can detect the MotC::His_6_ protein when expressed *in trans* on a multi-copy plasmid (Fig. 1C, lane 5). Furthermore, a strain expressing the MotC-His_6_ epitope tag variant on the chromosome is able to both swim and swarm comparable to the WT strain in plate-based assays (Fig. S1) indicating that the MotC-His_6_ protein is fully functional (compare *motC*::His_6_ images with those for the WT strain). The strains carrying the MotC-His_6_ or MotA-His_6_ MotC-His_6_ epitope tagged proteins also show motility indistinguishable from the WT (Fig. S1), indicating that the epitope tags do not interfere with the function of these proteins.

### All strains exhibit heterogeneous motility populations, each with a characteristic proportion of diffusive to superdiffusive cells

In a typical swarming motility assay, bacteria are transferred from liquid growth, and before collective swarming expansion, they find themself in a confined environment of soft agar (0.55% agar for this study) and a thin layer of liquid medium (Fig. 2A). This period is referred to as *swarming lag* (1), and little attention has been paid to this transition environment in which the cells become swarming competent. To investigate whether a strain with a MotAB, MotCD, or WT motor shows different single cell motility in this pre-swarming environment, we prepared miniature soft-agar plates using silicon spacers and inoculated a thin layer of liquid cell culture onto its surface (Fig. 2B). The microenvironment was sealed with a glass cover slip, incubated at 37°C and the motility of single cells tracked over a period of 8 hours.

**Figure 2.**
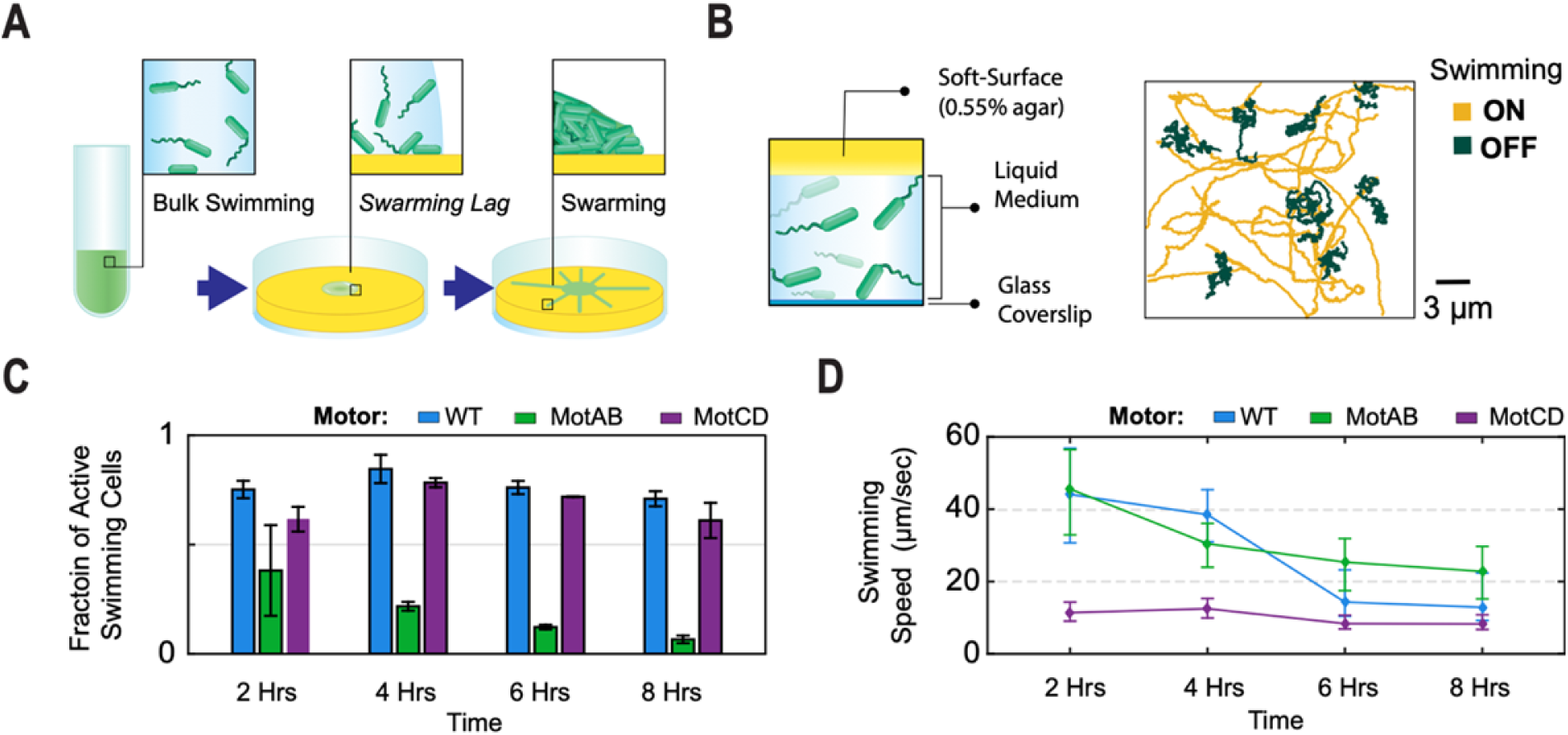
Measurement of fraction of active cells and their swimming speeds in the stagnant liquid-agar *swarming lag* phase environment. *(A)* Diagram of a swarming plate assay experiment. Three stages illustrated: (1) Inoculation of cells from liquid growth culture, (2) a *swarming lag* phase before cells reach confluency, and (3) the collective swarming expansion. *(B)* Illustration of experimental setup used for the *swarming lag* phase microenvironment (left). Two distinct populations were identified in this microenvironment setup (right): Cells with an active flagellum, ON, showed ballistic motion, while cells with an inactive flagellum, OFF, moved with a diffusive motion. *(C)* The fraction of active swimming population, ON, was quantified using the categories described in (B). The population activity was tracked every two hours for 3 min, over a period of 8 hours. Three replicates were used per strain; at least 140 trajectories were used per timepoint for each replicate. *(D)* Measured swimming speeds of the ON cells population in plot (C). At least 200 trajectories per timepoint. Error bars denote the first and third quartiles of the distribution about the mean.

We find that all strains show heterogeneously motile populations. We tracked cells and identified two populations with distinctive motilities: (1) flagellum-driven ballistic displacement trajectories with mean squared displacement, MSD, slope ≥ 1.4; these are referred to as swimming-ON cells (these trajectories are typical for active, propelled motion, rather than diffusive motion). MSD slope was calculated as defined previously (22). (2) trajectories with diffusive movement with MSD slope ≥ 0.7 but ≤ 1.3; we refer to these as swimming-OFF cells (Fig. 2B, right panel). Trajectories with MSD slope < 0.7 were considered as attached/semi-attached to the glass imaging surface and not used for this analysis. The fraction of swimming-ON cells was significantly larger for the WT and MotCD motor strains compared to the MotAB motor strain - about 2-fold after 2 hours (75.2%, 61.7% and 38.4%, respectively). This difference increased to 10-fold after 8 hours primarily due to a large drop in the fraction of swimming-ON cells among MotAB motor cells (6.7%); the fraction of swimming-ON cells remained relatively constant over time for WT and MotCD motor cells (Fig. 2C), with swimming-ON cells as a majority.

Although the MotAB motor strain had a small fraction of swimming-ON cells, the swimming speed was faster than the strain with the MotCD motor (45.6 ± 10.6 versus 11.4 ± 2.9 μ/sec at t=2hours; Fig. 2D). The MotAB motor strain progressively decreased its speed over time to 22.8 ± 7.3 μ/sec after 8 hours of incubation but remained significantly faster than the MotCD motor strain (8.3 ± 2.0 μ/sec). The WT motor started with a median speed of 44.1 ± 13.1 μ/sec after 2 hours, comparable to the MotAB motor strain; and it slowed down to 12.9 ± 6.6 μ/sec after 8 hours, closer to the MotCD motor strain speeds. The relatively constant diffusivity of the swimming-OFF cells indicates that the slowdown in swimming speeds across the strains over time is not due to an increase in viscosity under our experimental conditions (Fig. S2).

### Single cell tracking of swarming cells reveals MotAB and MotCD motors result in drastically different long-term intermittency in flagellar activity

To trace the origins of collective swarming motility by *P. aeruginosa*, it is important to not just measure population averaged behavior but to also determine the single cell behavior that may contribute to this collective behavior. To assess contributions of strains using one versus both stators during swarming motility, we tracked single cells from the edge of an early swarm—just as the tendrils begin to form. Cells harvested from the edge of the swarm were inoculated to the center of a miniature 0.55% agar plate (Fig. 3A). To track single cells in the crowded swarm environment, the initial swarming plates were inoculated with a co-culture containing a tracer fraction of cells carrying a constitutive green-fluorescence protein (GFP) plasmid (Fig. 3A) at 1-5% (v/v) of GFP cells.

**Figure 3.**
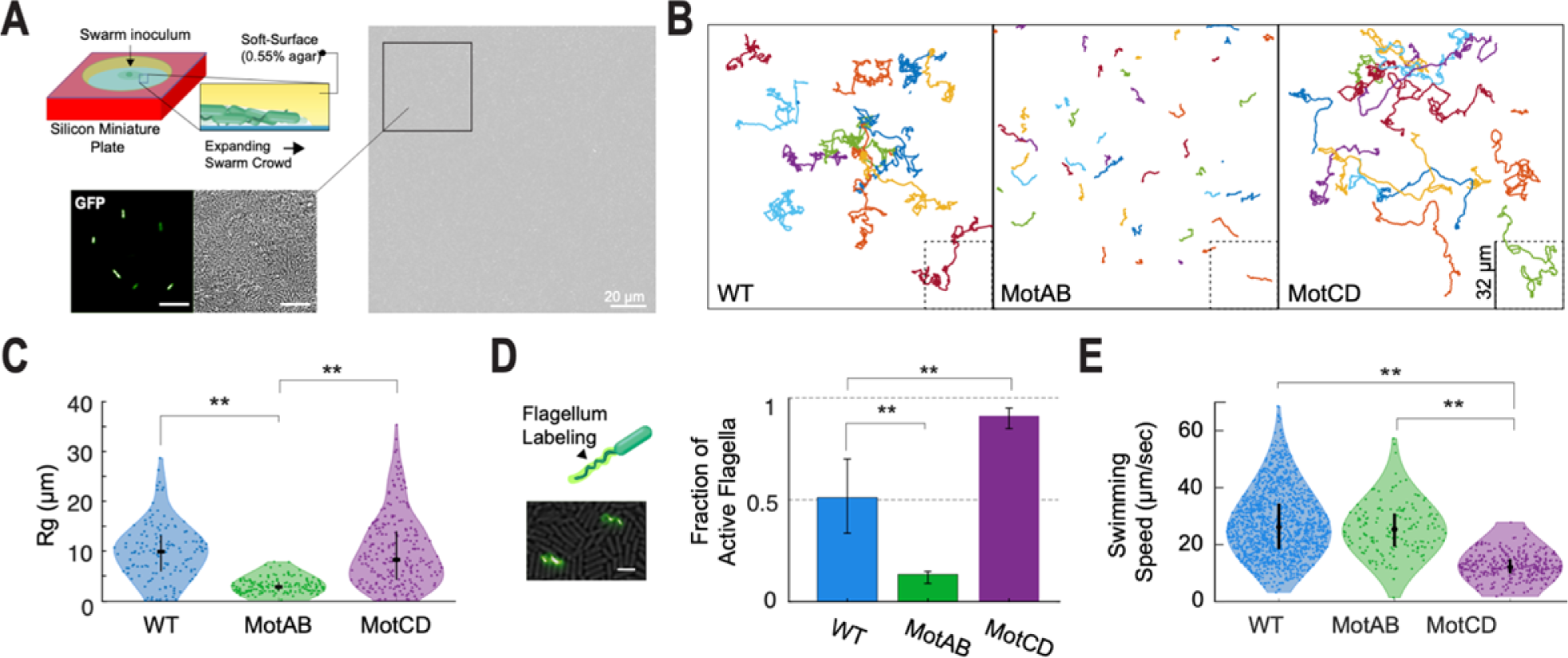
Single-cell motility measurement of cells in a crowded environment of their own kin and 2D confinement. *(A)* Diagram of experimental setup for tracking swarming bacteria on a soft-agar medium surface (top left). Cells harvested from a swarm plate with 1-5% (v/v) co-culture of cells carrying a constitutively expressed copy of GFP on a plasmid (lower left, right). This approach permitted precise single cell tracking in an environment crowded with thousands of cells per field-of-view. 10μm scale-bar in zoom-in inserts. *(B)* Representative trajectories of tracked cells in the crowded environment over a period of 30 seconds. The trajectories presented for each indicated strain come from a compilation of different fields of views from at least 3 replicates. *(C)* Violin plot of measured radius of gyration for the single cells trajectories in the crowded environment for the three indicated strains, as displayed in *(B)*. At least 139 cells were tracked per strain. *(D)* Co-inoculation containing a small fraction of cells, 5-20% (v/v), with a FliC^T394C^ mutation for maleimide staining was used for direct quantification of actively rotating flagella. The bar plot on the right reports the fraction of active flagella observed for each strain. At least 12 fields-of-views from four replicates were used and at least 200 flagella per strain were counted. 5μm scale-bar. *(E)* Swimming speeds of cells moving in a 2D thin liquid volume medium confined between a 0.55% agar surface and imaging glass coverslip, under a diluted cell volume fraction, Φ (Movie S1). At least 180 trajectories were measured per strain. For panels C-E data sets were analysed by one-way ANOVA followed by Tukey’s post-test comparison. \**P* < 0.05, \*\**P* < 0.001; ns, not significantly different.

In this crowded condition, the MotAB motor strain showed significantly shorter translational displacement compared to WT and MotCD motor strains (Fig. 3B). Such differences are quantitatively reflected by the radius of gyration (Rg) of the trajectories, which describe a characteristic spatial extent of cell travel at the longest observation time (23). The Rg is defined as:

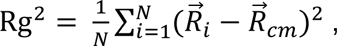

where *N* is the number of points in the track, 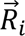 is the position vector of the *i^th^* point on the trajectory, and 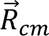 is the center-of-mass of all points. The WT and MotCD motor strain display trajectories up to 10 times longer than the mean trajectories of the MotAB motor strain (Rg = 6.2 ± 3.8, 6.1 ± 4.5 μm and 1.9 ± 0.9 μm, respectively; Fig. 3C).

To quantify the flagellar activity that drives the motility behaviors observed under the crowded conditions described above, we harvested cells from the edge of an early swarm from a co-culture containing a tracer population of cells carrying a threonine-to-cysteine mutation in the flagellum filament subunit (FliC^T394C^) (24), analogous to the single cell tracking approach described above in Figure 3A (with 5-20% v/v of FliC^T394C^ cells). The flagella were stained with an Alexa Fluor 488 C5 maleimide added to the plate prior to harvesting (Fig. 3D). This approach allowed for direct observation of the fraction of active flagella. While almost all the flagellated cells using the MotCD motor were active (92 ± 6%) only 13 ± 9% of the MotAB motor strain had an active flagellum (Fig. 3D, Movies S1-2). The WT motor had an intermediate fraction of active flagella, but also with the largest variation, 51 ± 24% (Movies S3). The percentage of active flagella observed here aligns with the differential percentage of motile cells measured above during *swarming lag*.

Finally, we characterize the flagellar output under this confined monolayer geometry. Since in the crowded regime is not possible to untangle the flagellar output from the contributions of all neighboring particles to the movement of a single cell, we diluted the system to measure the free-swimming speeds of cells in a 2-D confined (monolayer) liquid volume. Cells harvested from the edge of a swarming motility plate were diluted and inoculated as a thin liquid film, ∼1.5 μm in height, between a 0.55% agar surface and an imaging cover glass (Movie S4). In this free-swimming configuration, the WT and MotAB motor outperform the MotCD motor by about 2-fold (26.2 ± 11.5, 25.4 ± 8.8 and 12.1 ± 4.2 μm/sec, respectively). Hence, the agar-liquid-glass monolayer configuration imposes a characteristic load on the flagellum equivalent to a moderate viscosity, i.e., ∼10 cP (Fig. 1B). Therefore, under this 2-D monolayer configuration, as crowding increases (Fig. 3A), we expect such flagellar load to increase due to cell-to-cell interactions (collisions) and lead to a convergence in motor output between the strains like the measured behavior above with increasing flagellar viscosity load.

### Modeling of cell populations with heterogeneous motor output reveals the existence of unanticipated unjamming transition modalities

To examine how diverse flagellar motor outputs and intermittency (i.e., fraction of flagellum-ON cells) in a heterogeneous population are integrated into either a collective swarming or non-swarming phenotype, we designed a physical simulation model of self-propelled rods (aspect ratio = 4) in a 2D crowded environment with a volume fraction, Φ, of 0.96. These are conditions akin to the high-density environment of swarming cells. Each simulation contained a fixed fraction of flagellum-OFF (*F_f_*=0) and flagellum-ON cells (*F_f_*>0) (Fig. 4A, Movie S5-8). The relative magnitude of the active force is reported compared to the repulsive interaction coefficient, k, of the repulsive linear spring potential for the particles (see SI Appendix). To evaluate the extent of movement within the crowd, we report here the collective radius of gyration, Rg, for the systems normalized by the radius of gyration of the homogeneous system with all flagellum-OFF particles and denoted as Rg_N_ (Fig. 4B).

**Figure 4.**
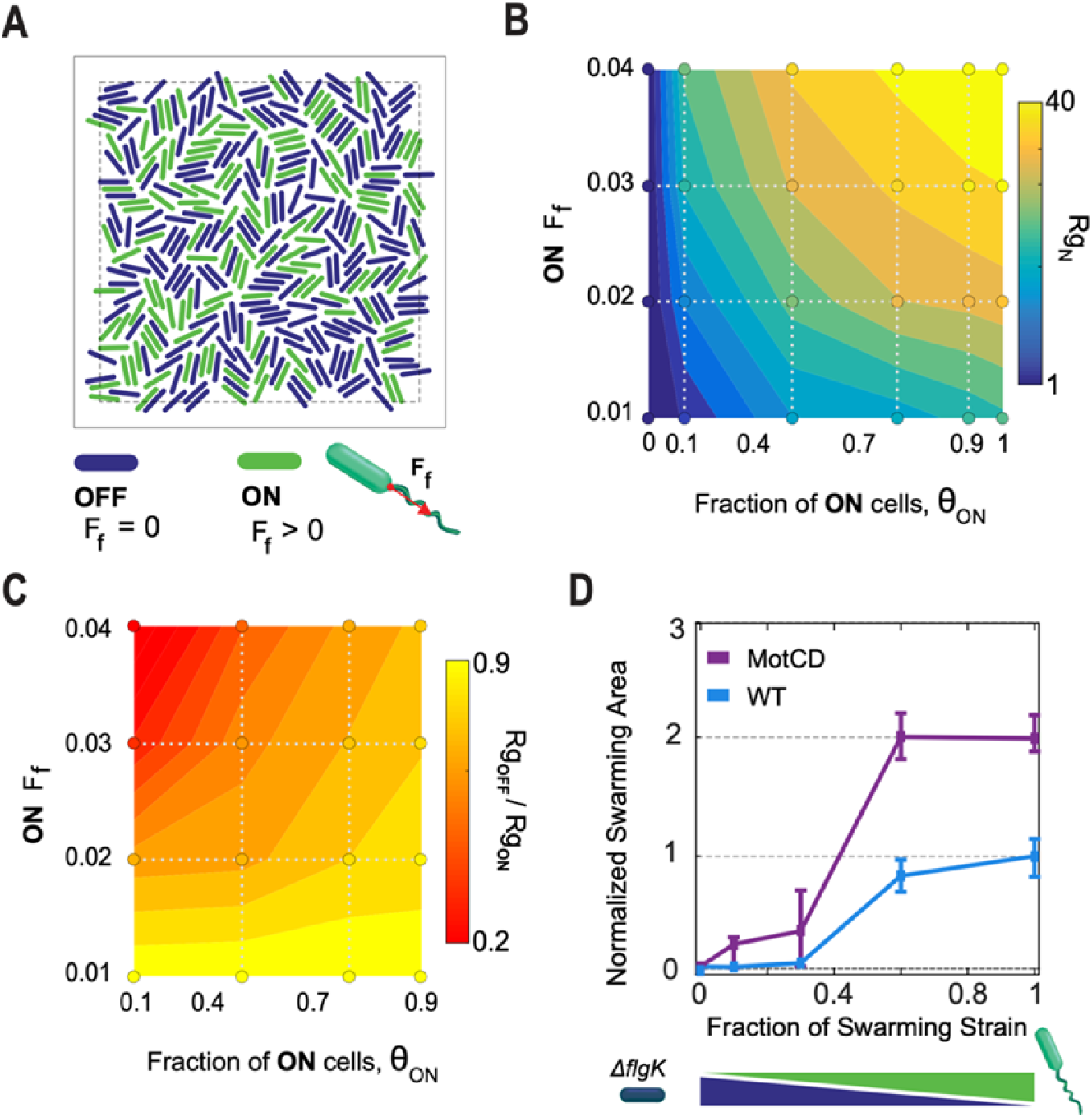
Physical modeling of the crowded environment predicts a landscape of unjamming transitions for different combinations of the flagellum motor force output and fraction of active flagellum cells. *(A)* Simulations of a crowd of self-propelled rods to evaluate the influence of population heterogeneity, ON-vs OFF-flagellum, and varying flagellar force outputs on collective motility. The simulations were tested at a volume fraction (Φ) of 0.96, and cell aspect ratio of 4. The fraction of ON cells (θ_0*N*_) and their flagellar output (F_f_) were varied. *(B)* Normalized mean radius of gyration, Rg_N_, for the particles in the tested crowded systems as illustrated in *(A)*. The values were normalized by the mean radius of gyration of the homogeneous system with all flagellum-OFF particles (F_f_=0). Contour map was estimated by interpolation between the grid of tested conditions (circular markers). *(C)* The asymmetry in translational movement in the heterogenous systems between the flagellum-ON and -OFF particles was measured by the ratio in mean radius of gyration for the two populations (Rg_OFF_/Rg_ON_). A Rg_OFF_/Rg_ON_ of 1 corresponds to equal translation by both particle types in the heterogenous crowd. *(D)* Normalized swarming area as a function of increased concentration of the swarming strain (WT and MotCD motor) in mixed culture with the flagellum deficient Δ*flgK* mutant. The Δ*flgK* strain lacks a functional flagellum, and hence is swarming deficient. Error bars denotes the quartiles of the distribution about the mean.

The progressive increase in fraction of flagellum-ON cells promotes incremental movement via a dynamical and cooperative transition analogous to an unjamming transition in the field of active fluids (25–28). For homogeneous systems of flagellum-ON cells, translation is positively dependent on the flagellar force, with a Rg_N_ of 9.28 when *F_f_*=0.01 to up to a Rg_N_ of 68.67 for *F_f_*=0.04. The data shows that it is possible to achieve comparable, large displacements under multiple configurations. For instance, a system with a 0.9 fraction of flagellum-ON cells (θ_ON_) and *F_f_* = 0.02, and a system with θ_ON_ = 0.4 and *F_f_*=0.03 will promote similar crowd movement, with Rg_N_ of 24.1 and 22.6 respectively (Fig. 4B). However, the systems motility is not homogeneously shared by the flagellum-ON and flagellum-OFF particles, as reflected by their Rg_N_ ratio between the two population types, Rg_OFF_/Rg_ON_. For example, while a system with [θ_ON_ = 0.9, *F_f_* = 0.01] and a system with [θ_ON_ = 0.1, *F_f_* = 0.03] achieve a similar Rg_N_ of about 7.5, the asymmetric translation between the flagellum-OFF vs -ON is greater for the latter system with a Rg_OFF_/Rg_ON_ ratio of 0.31 compared to a 0.83 Rg_OFF_/Rg_ON_ ratio for the first system (Fig. 4C). Hence, in the first configuration, the flagellum-ON population creates a more uniform cooperative collective movement, i.e., close to Rg_OFF_/Rg_ON_ = 1, that supports the flagellum-OFF population movement; on the other hand, system with lower Rg_OFF_/Rg_ON_ ratio creates a situation in which the flagellum-ON population moves through the flagellum-OFF population. A cognate behavior has been observed in heterogenous mixtures of hyperswarming and swarming strains (29), or species (30), in which the hyperswarming cells move through the swarming strain to lead the swarming front. Therefore, in heterogenous systems, like bacterial populations, the interplay between *F_f_* and θ_ON_ may affect *cooperative* vs *selfish* motility between the diverse members of the population.

### Modulation of swarming motility in *P. aeruginosa* via modulation of flagellum-active populations

Based on the observed characteristic dynamic distribution of inactive cells between the WT, MotAB and MotCD motor strains and our model estimations on how such active to inactive ratios will impact the promotion or arrest of swarming motility, we predict that altering the proportion of inactive cells in a population of swarm-competent cells would impact swarming behavior at a macroscopic level. To test if and how the fraction of inactive cells impacts swarming, we controlled the fraction of active cells by mixing the swarming strains (WT and MotCD motor) with a flagellum-less strain (Δ*flgK*) at different static ratios, similar to an approach we used previously (31). As shown in Fig. 4D, swarming motility of both the WT and MotCD motor strains is negatively impacted by increasing the proportion of inactive Δ*flgK* cells in the population, consistent with our prediction and our previous report (31). For the MotCD strain, swarming onset occurs when MotCD cells comprise ∼0.1 to 0.3 fraction of the mixed population. To calculate the fraction of cells with active flagella in this mixed population, we must consider our results above (Fig. 3D), whereby the WT and MotCD swarming strains exhibit distinct proportions of flagellum-active and -inactive cells in their populations. Taking such data into account, the fraction of the MotCD strain with active flagella at the onset of swarming in the MotCD/Δ*flgK* mixed population is ∼ 0.18 (i.e., 0.9 fraction of active cells in the MotCD strain alone and an average of 0.2 fraction of the mixed MotCD/Δ*flgK* population at swarming onset). For the WT strain, the fraction of the WT population with active flagella at the onset of swarming is ∼ 0.15 (with 0.5 fraction active cells in WT and 0.3 fraction of the mixed WT/Δ*flgK* population). Notably, both values are comparable to the measured fraction of flagellum-active cells of the swarming deficient strain, MotAB motor (∼13%), indicating that increasing the fraction of inactive cells in WT or MotCD motor swarming populations can effectively mimic the lack of swarming observed for the MotAB motor strain.

### Pro-flagellar shutdown effect of MotAB stator in WT can be offset by an increase in MotCD production

Given that expression of MotAB dominates that of MotCD, we hypothesize that the MotAB stator maintains motor recruitment in the WT motor during swarming, hampering the heterogenous motor from reaching higher degrees of swarming, e.g., MotCD motor-like hyper swarming. A strong prediction of this model is that a MotCD-dominated motor would enhance swarming motility, a prediction that aligns with the data shown in Fig. 4D. We previously showed that deleting the MotAB stator results in a hyper-swarming phenotype (15, 20). Here, we utilized a plasmid carrying the *motCD* genes, whose expression is under arabinose-inducible control, to increase expression of MotCD in a WT background. We observed an arabinose-dependent increase of swarming motility with increased expression of MotCD (Fig. 5A). At 0% arabinose, the swarming motility between the WT pMotCD strain and the WT carrying the empty vector (pMQ72) was not significantly different (Fig. S4A). When induced MotCD expression (1% arabinose) led to a mean increase of ∼40% in swarming area compared to the WT pMQ72. In contrast to its significant impact on swarming, increased expression of the *motCD* genes did not affect the swim speed distributions (Fig. S5), a finding consistent with the data shown in Fig. 1B and with a role for MotCD function specific to the high viscosity environment associated with swarming motility.

**Figure 5.**
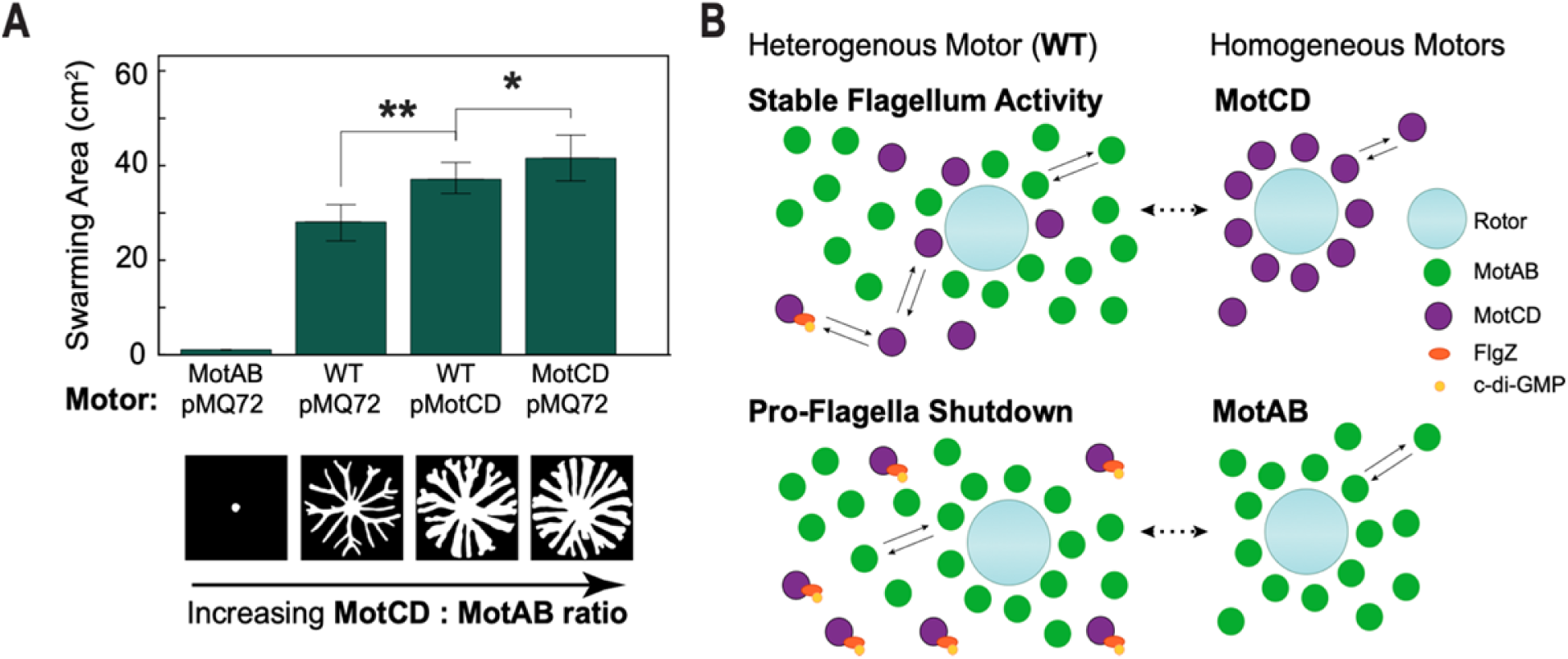
Increasing MotCD:MotAB ratio leads to increased swarming motility. *(A)* Expression of MotCD via an arabinose-inducible plasmid (pMotCD) increases WT swarming phenotype (pMQ72 is an empty vector control). All strains were grown in soft agar swarming plates with 1% arabinose. No significant difference was observed between the WT/pMQ72 and WT/pMotCD at 0% arabinose (Fig. S4A). At least 6 plate replicates per condition. Error bars denote the first and third quartiles of the distribution about the mean. \**P* < 0.05, \*\**P* < 0.001; ns, not significantly different. Data were analysed by one-way ANOVA followed by Tukey’s post-test comparison. Lower panel shows representative segmented swarm areas for the four different stator ratio configurations. *(B)* Model of stator type dynamics and their expected influence on the motor intermittency in a crowded swarming environment (high flagellar load). All three motor types are expected to maximize flagellar output under this condition, i.e., motors fully, or mostly, decorated with stators (Fig. 1B & 3E). The heterogenous WT motor asymmetrically recruits MotAB stators due to its higher affinity at low-to-mid range viscosities and its elevated expression compared to the MotCD stator (Fig. 1B-C). To further hinder the recruitment of MotCD, we previously described the cyclic di-GMP-dependent binding of FlgZ to the MotC subunit of this stator and its sequestering from the motor (15, 46). We postulate here that like the MotCD homogenous motor, the presence of MotCD stators helps stabilize the flagellum activity, i.e., maintaining an active flagellar motor. The MotCD motor sustains population with 92% active flagella (Fig. 3D). In contrast, the MotAB homogenous motor is prone to a flagellar shutdown with about 88% of its population having an inactive flagellum (Fig. 3D). As well as modulating the flagellar torque output, the MotAB and MotCD stators may integrate molecular signals to regulate the flagellar activity state, long-term intermittency, among the heterogenous population.

## DISCUSSION

In this work, we propose a generalizable conceptual framework through which the collective flagellum driven motility in a population of *P. aeruginosa* cells can be controlled via dynamic MotAB/MotCD stator recruitment. An important unanticipated observation presented here from direct measurement of single-cell flagella tracking in a crowd of swarming cells is that MotAB and MotCD motors have drastically different intermittency in flagellum activity (∼12% MotAB versus ∼92% of MotCD motors are flagellum active) (Fig. 3D). This finding implies that stator recruitment can strongly impact the fraction of time that a cell’s flagellum is on. The implications of this observation are broad, as outlined below.

It is possible to rationalize the behavior of heterogeneous motile populations of *P. aeruginosa* cells with different stator recruitment in their flagellar motors and different resultant activity levels by importing concepts from studies of ‘active granular matter’ (32–35). Based on experimental and modeling data, we propose that swarming occurs when numerous individual motility choices achieved via stator recruitment are successfully integrated into macroscopic community motion via processes related to the ‘unjamming’ transition, which separates a static solid-like biofilm from a flowing liquid-like swarm in ‘granular matter’, such as floes in drift ice that cause jams in freshwater rivers. For example, instead of a single value of flagellum-generated force or active fraction, we find that a sliding scale of different combinations of these parameters can all achieve community motility. A group of sufficiently crowded particles are prevented from flowing and from exploring possible configurations in phase space. From this perspective, it is possible that *P. aeruginosa* can swarm readily as a monotrichous bacterium because it can generate diverse combinations of motility behaviors via stator recruitment: even cells that are slower or not actively pushing can, counterintuitively, promote swarming of the population by effectively creating local free volume and allowing flow. In a more general compass, these results in *P. aeruginosa* suggest that deployment of temporal heterogeneity in a motile microbial population can drive unexpected collective motility behavior.

Swarming has traditionally been described by requirements that are deterministic in nature, in terms of necessary and sufficient molecular components (e.g., the appropriate stators). The data here indicate that this perspective is too reductionist: Not only are single cell motilities heterogeneous, but they are also characterized by multimodal distributions in space and time, with shifting subpopulations (Fig. 2C-D). We find that populations with either MotAB, MotCD, or WT motors, *all* exhibit heterogeneous motility populations, each with distinct characteristic proportions of highly motile superdiffusive cells. We propose that a homogenous MotAB motor inhibits flagellar activity by decreasing the fraction of time this motility appendage is active, while the presence of MotCD stators in the motor leads to a steadily active flagellum motor (e.g., the WT and MotCD motors) (Fig. 5B). We propose that this effect of MotAB-induced flagellum intermittency may be the first step in a full flagellum shutdown in high viscosity environments, i.e., high flagellar load. Indeed, our proposed picture is not inconsistent with previous experimental observations in which only ∼3.5% of flagellum-tethered MotAB motors spin (13). In fact, we find that a MotAB motor is especially prone to shutdown due to intrinsic intermittent activity, which we observe even in cells not associated with the surface.

The concept of a *P. aeruginosa* population that is structurally heterogeneous because it is capable of adaptive stator recruitment allows us to engage recent results on heterogeneity from a new perspective. We have shown that a population with heterogeneous motility can integrate into a swarm via different combinations of parameters, such as the fraction of active flagellum cells or the flagellum motor force output (Fig. 4B). Our simulations and experimental data suggest that having too small a fraction of flagellum active cells can lead to swarming arrest, even if mechanically, each cell has torque output comparable to other swarmers, e.g., the swarming deficient strain that uses MotAB stators. Although increasing the fraction of flagellum active cell is a requirement for increased mobility of crowded systems, a majority of flagellum active cells is not necessary. Even when the flagellum inactive cells are the dominant population, the system may be able to maintain motility, as seen in the WT and MotCD motor strain swarming inhibition experiments (Fig. 3D), as well as recent studies by Hogan et al. (31) and Xavier et al. (36, 37).

During swarming, instead of a paradigm where MotAB contributes less to swarming than MotCD, we find that MotAB and MotCD have in fact rivaling contributions to flagellar activity (or inactivity) during swarming, with MotCD maintaining an actively motile population to facilitate swarming motility and MotAB maintaining instead a population with individual cells that exhibit infrequent spasms of motility. Indeed, flagellar population intermittencies may have been already optimized by other swarming bacteria species through evolution as most of these express multiple flagella (1). Although *P. aeruginosa* manages to swarm with a single flagellum, such collective motion is dramatically enhanced with the expression of multiple flagella (29). Given the similar flagellar output of single- and multi-flagellated *P. aeruginosa* (38), we expect the increased swarming motility to arise from an increase of flagellar population intermittency.

We note that different regulators of motility, such as cell-cell pili interactions (31, 39), secretion of extracellular polymeric substances (EPS) (31, 40), and secretion of rhamnolipids (36), have all been observed to impact swarming. However, from the present perspective, it is interesting to see how they relate to sensing events that may parallel swarming. For example, how do bacteria sense local effective viscosities or the ‘crowdedness’ of local environments? These molecular factors alone can affect the rheology of the media that the bacteria encounters (41–43), however in a swarm bacterium packing fraction may also change the effective viscosity, or flagellar load, that a single cell experiences. There are numerous examples in the colloidal suspension literature (34, 44) showing that the effective viscosity that the suspended particles experience increases dramatically relative to the nominal viscosity of the solvent as the packing fraction of the colloids increases toward the onset of jamming (35), which arrests collective movement. A similar effect occurs in the crowded intracellular environments which have macromolecular packing fractions approaching 30-40%. One cannot accurately estimate diffusion times of proteins and other metabolites from the viscosity of water since diffusive transport is dramatically slowed by the presence of macromolecules (45).

Since the two stator types apparently have different viscosity sensitivity for motor recruitment, shown both here and elsewhere (5), it raises the question of how *P. aeruginosa* measures viscosity or crowding. If the bacteria can measure viscosity by how much torque is necessary to achieve a given speed, or sense collisions with other bacteria, their motor can be sensitive to the effective viscosity, or effective flagellar load, and not the background viscosity of the fluid. This result would provide a mechanism for the MotCD stator to become the predominant motility stator in a dense, swarming environment, even though the background viscosity may favor the MotAB stator. Such asymmetric preference may nonetheless be impacted by stator availability (Fig. 1C) as well as intracellular cyclic di-GMP (c-di-GMP) levels (20).

Previously we showed that PilZ domain-containing protein FlgZ binds to MotC in a c-di-GMP dependent manner which can prevent the MotCD stator integrating into the flagellum motor (15, 46). Such a motor state combined with heavy flagellar loads would render the resulting MotAB motor effectively inactive, since this stator is prone to infrequent flagellum activity or even flagellum shutdown (Fig. 5B) (47). Strains that exhibit flagellar shutdown or flagellum impairment are often linked to increases in EPS production (48). Therefore, a population with inactive flagella may produce the EPS that further antagonizes swarming (31, 46, 49). We speculate that in addition to its ability to act as a ‘flagellar dynamometer’ (50), flagellar load sensing when combined with other factors such as cell-cell (51) or surface interaction (13, 52–54) may influence the motor activity. Our findings, combined with the recent studies noted above, are critical for future model building of swarming motility control.

The conceptual results presented here are in principle generalizable. Beyond an explication of flagellum-driven swarming phenomena, a collective shutdown of flagellum motility orchestrated by stator recruitment may be important to surface sensing mediated signaling pathways that lead to flagellum shutdown and nucleation of microcolonies in heterogeneous bacterial populations for biofilm formation.

## Materials and Methods

### List of Strains

All strains are listed in SI Appendix Table S1. Plasmid pSMC21 constitutive expresses *gfp* and confers resistance to kanamycin (200 μg/mL). Plasmid pMQ72 allows for inducible expression by addition of arabinose and confers resistance to gentamycin (25 μg/mL).

#### Strain construction

In-frame insertion of the His_6_ epitope tag into the *motA* and *motC* genes was performed via allelic exchange, as previously described (55). Plasmids for this purpose were constructed via cloning by homologous recombination of relevant PCR products into the pMQ30 vector using Gibson assembly. Constructs for plasmid-based expression of genes were generated using PCR and Gibson Assembly^®^ (NEB, Boston, MA) followed by cloning into pMQ72. For all plasmids and constructs used in the experiments described herein, the relevant cloned genes were fully sequenced to confirm that the correct sequences were present. PCR and sequencing was also used to confirm the presence of the His_6_ epitope tag in the *motC* gene on the chromosome in the *motA*::His_6_ *motC*::His_6_ and *motC*::His_6_ strains.

#### Protein detection and cellular localization experiments

Bacterial strains were grown either in liquid cultures or on swarm agar plates in M63 minimal salts medium supplemented with 1 mM MgSO_4_, 0.2% glucose and 0.5% CAA with 0.5% arabinose for induction of plasmid-based expression of the pMotC::His_6_ protein. Whole cell lysates and membrane fractions were prepared as previously described (56). Total protein concentrations in membrane fractions were quantified using the Pierce™ BCA protein assay kit (ThermoFisher Scientific, Waltham, MA). For Western blotting, equivalent total protein quantities from membrane samples were resolved by SDS-PAGE using Any kD^TM^ polyacrylamide gels (Bio-Rad, Hercules, CA). Proteins transferred to a nitrocellulose membrane were probed with a monoclonal anti-His_6_ antibody (Qiagen, Germantown, Maryland). Detection of proteins via Western blotting was performed by fluorescence detection using IR-Dye^®^-labeled fluorescent secondary antibodies and imaged using the Odyssey CLx Imager (LICOR Biosciences, Inc., Lincoln, NE). Protein quantification was performed using Image Studio Lite software (LICOR Biosciences, Inc., Lincoln, NE).

#### Motility assays

Swarm motility plates were prepared with M63 medium supplemented with 1 mM MgSO_4_, 0.2% glucose and 0.5% CAA and 0.05 % arabinose, referred to as M63 medium in this report, and solidified with 0.5% agar. Swarm assays were performed as previously described (57). Swim motility plates were prepared with M63 medium supplemented with 1 mM MgSO_4_, 0.2% glucose and 0.5% CAA and solidified with 0.3 % agar. Swim assays were performed as previously described (58).

#### Single-cell swimming motility tracking

Cell tracking was performed as previously described (24) with a few minor adaptations. *P. aeruginosa* PA14 strains were incubated in liquid LB medium with shaking at 37°C overnight. Cells were washed with M63 medium; for the strains crying an arabinose inducible plasmid pMQ72, the M63 medium was supplemented with 1% arabinose. The washed culture was diluted to OD_600_ ∼1.0; 20μl of this culture were inoculated into 1mL of M63 medium at different viscosities and incubated for 1.5 hrs without shaking at 37°C before imaging. Methylcellulose 400cP powder (Sigma-Aldrich) at varying percent concentrations was used to change the viscosity of the M63 medium following the manufacturer directions for methylcellulose hydration. Glucose (0.2%), MgSO_4_ (1mM), and Casamino Acids (0.5%) after hydration.

The cells were injected into a flow cell channel (Ibidi sticky-Slide VI0.4 with a glass coverslip). Bright-field imaging recordings were taken in using a Phantom V12.1 high speed camera (Vision Research) with a 5 ms exposure at 200 frames per second (fps) and a 0.1 μm/pixel resolution on 600 x 800 pixels field-of-view (FOV). The imaging protocol was performed on an Olympus IX83 inverted microscope equipped with a 100x oil objective, a 2x multiplier lens, and Zero Drift Correction autofocus system and a heating stage (30°C).

Image processing and cell-tracking of the near-surface swimming recordings were processed with algorithms written in MATLAB R2015a (Mathworks) described in previous work (24). Swimming trajectories were identified as those that traced a radius of gyration ≥ 2.5 μm and a ballistic movement with MSD slope ≥ 1.4 these thresholds discriminated against too short and/or passive (diffusive) trajectories. The reported speeds per trajectory are the calculated median speed from a distribution displacements using 20 frames (100 ms) moving window over the full trajectory.

#### Agar-liquid film cultures

To mimic the *swarming lag* phase of a swarm plate assay, warm 0.55% agar M63 medium was poured onto a 25×25×2.5 mm silicone CoverWell^TM^ imaging chamber (GraceBio) with a bottom glass coverslip. Immediately after, a glass coverslip was laid to create a flat surface. After 1h, the top coverslip was removed, leaving a flat soft agar surface. The surface was allowed to dry for ∼20 min until a slight meniscus is formed. 10μl of washed and diluted (to OD_600_ ∼1.0) overnight culture were inoculated onto the agar surface, as described above. A clean glass coverslip was laid on top of the inoculated culture to create a thin film of liquid culture sealed between agar and the glass imaging surface. The sealed imaging chambers were incubated at 37°C and only taken out of the incubator for imaging, ∼10 min, and returned to incubator. Imaging and data processing was performed as above for single-cell swimming motility tracking. Trajectories were classified as: flagellum ON, a ballistic movement with MSD slope ≥ 1.4; or flagellum OFF, a diffusive movement with MSD slope ≥ 0.7, but ≤ 1.3. Cells with MSD slope < 0.7 were considered surface attached. For flagellum ON trajectories, the reported speeds per trajectory are the calculated median speed from a distribution displacements using 20 frames (100 ms) moving window over the full trajectory.

#### Crowded environment assays

A 25×25×2.5 mm silicone CoverWell^TM^ imaging chamber (GraceBio) with a bottom glass coverslip was filled with heated M63 medium containing 0.55% agar; and glass coverslip was pressed on top, as above. The top coverslip was removed, and excess liquid was allowed to dry, ∼5 min. To inoculate the miniature plates, cells were harvested from a swarming motility plate–after 10 hrs incubation. Using a toothpick to collect a sample from the tip of a tendril, or edge of the colony for swarming deficient strains, the sampled biomass was placed at the center of the agar surface on the miniature plate. A clean imaging coverslip was gently pressed on the top to firmly enclose the cells against the agar and conserve humidity; smearing or distortion of the inoculum from the center as carefully avoided. The miniature plates were incubated at 37°C for 3h. After incubation period, the cells traveled radially outwards from the center inoculum and imaged.

To track individual cells in the expanding crowded environments, a fraction of cells constitutively expressing a GFP-carrying plasmid, pSMC21, were co-inoculated in the source swarming motility plate. The strains with and without the GFP reporter were mixed to a 1:99 volume ratio, respectively, from separate washed and diluted (OD_600_ ∼1.0) overnight cultures. The co-culture was inoculated to a swarming motility plate and incubated as before. The plate was sampled after 10 hrs incubation as described above. A higher volume ratio of 5:95 was used for swarming deficient strains due to their lower collective motility in the confined crowded environment which reduces their level of mixing as the crowd expands during incubation in the miniature agar plate, i.e., only the ΔMotCD pSMC21 cells closest to the edge of the new sampled inoculum will be carried by the expanding front. No significant changes in swarming phenotype were observed for the strains carrying the plasmid pSMC21 (Fig. S3) compared to the background strain.

Imaging recordings were taken with an Andor Neo sCMOS camera with Andor IQ software on an Olympus IX83 microscope equipped with a 100x oil objective and Zero Drift Correction 2 continuous autofocus system. To avoid bias of the constrained edge cells, the XYZ location for the recording was set ∼15μm from the edge of the expanding crowded environment. One minute fluorescence recordings were taken with 100 ms exposure for a 10fps recording (shutters were continuously open and without display feedback to maximize frame rate) with Lambda LS (Sutter Instrument) xenon arc lamp and a green fluorescent protein (GFP) filter. The image size was 133 μm by 133 μm (2048 by 2048 pixels). Except for Δ*motCD* strain, all other strains were imaged with this protocol. The Δ*motCD* strain was imaged with 100 ms exposure at 1 fps for 1 min with active shutter to minimize accumulation exposure; this strain was more sensitive to fluorescence exposure. Image processing and cell-tracking of the GFP fluorescent cells were processed with algorithms written in MATLAB R2015a (Mathworks) as described above.

Individual flagella tracking was performed similarly as above. A *P. aeruginosa* PA14 strain with a FliC protein modified, a threonine-to-cystine mutation, FliC^T394C^, as previously described (24), was co-inoculated with a strain not carrying this mutation. The strains with and without the FliC^T394C^ mutation were mixed to a 5:95 volume ratio, respectively, from separate washed and diluted (OD_600_ ∼1.0) overnight cultures; similar to the single cell tracking in a crowded environment described above, higher volume ratio of 1:4 was used for swarming deficient strains. The mixed culture was inoculated to a swarming motility plate and incubated as before. After incubation, the flagella were stained using Alexa Fluor 488 C5 Maleimide (Molecular Probes) at 10 μg/mL. The edge of a tendril, or colony, was stained with ∼1 μL of the stock stain and allowed to sit for 10 min before the biomass was sampled for imaging. Imaging protocol was the same as described above. Only intensity rescaling was used on the raw fluorescent images. Quantification of active and inactive flagellum was done manually using MATLAB R2015a to display the recording.

#### Free-swimming on 2D-agar surface

A miniature soft agar plate was prepared as described above. After removal of top glass coverslip, 2μL of M63 media was placed on the agar surface. Cells were sampled from a standard swarm plate using a plastic inoculation loop. The loop was smeared on a fresh swarming motility plate to dilute the sample. The diluted sample on inoculation loop was gently caressed over miniature agar plate added liquid medium. An imaging glass coverslip was gently pressed to seal the silicon agar surface for imaging. Only flat areas where cells were visibly swimming in a constrained 2D space of media (∼1-2 μm of space between agar surface and glass coverslip) were used for this measurement. Bright-field imaging using a high-speed camera (Vision Research), image processing and single-cell tracking was carried as described above.

#### Mixed population swarming assays

Swarming assays of mixed swarming deficient strains (*ΔflgK* – lacking a flagellum) and the different swarming strains were performed as described above, with some modifications. Overnight cultures of the selected strains were washed using M63 medium and then normalized to an OD_600_ of ∼1.0. The washed cultures were then mixed to the desired testing ratios to a final volume of 100 μL. Finally, 2 μL of the co-culture mixture was inoculated to a 0.55% agars swarm plate assay as described before and allowed to grow for 18 hours at 37 C. Plates where not stacked to avoid temperature gradients on the plates.

#### Statistical analysis

One-way ANOVA with multiple comparisons was performed pairwise between all isolates using the GraphPad Prism 6 software or Matlab software.

## Acknowledgements

This work was supported by NIH RO1 R01AI143730 to G.C.L.W. and C.K.L., NIH R37 AI052453 to G.A.O. and S.S.W. and NSF PoLS 2102789 to C.S.O. J.D.A. is supported by NSF Graduate Research Fellowship Program DGE-1650604. We received support from the Bio-MT Molecular Tools and Molecular Interactions and Imaging Core (P20-GM113132) at the Geisel School of Medicine at Dartmouth.

## References

1. D. B. Kearns, A field guide to bacterial swarming motility. Nature Reviews Microbiology 8, 634–644 (2010).

2. J. Yan, H. Monaco, J. B. Xavier, The ultimate guide to bacterial swarming: an experimental model to study the evolution of cooperative behavior. Annu Rev Microbiol 73, 293–312 (2019).

3. Y. Sowa, R. M. Berry, Bacterial flagellar motor. Quarterly Reviews of Biophysics 41, 103–132 (2008).

4. A. Paulick et al., Dual stator dynamics in the Shewanella oneidensis MR-1 flagellar motor. Mol Microbiol 96, 993–1001 (2015).

5. Z. Wu, M. Tian, R. Zhang, J. Yuan, Dynamics of the two stator systems in the flagellar motor of *Pseudomonas aeruginosa* studied by a bead assay. Applied and Environmental Microbiology 87, e01674–01621 (2021).

6. M. Beeby et al., Diverse high-torque bacterial flagellar motors assemble wider stator rings using a conserved protein scaffold. Proceedings of the National Academy of Sciences 113, E1917–E1926 (2016).

7. M. Kaplan et al., The presence and absence of periplasmic rings in bacterial flagellar motors correlates with stator type. eLife 8, e43487 (2019).

8. J. Haiko, B. Westerlund-Wikström, The role of the bacterial flagellum in adhesion and virulence. Biology (Basel) 2, 1242–1267 (2013).

9. S. W. Reid et al., The maximum number of torque-generating units in the flagellar motor of *Escherichia coli* is at least 11. Proceedings of the National Academy of Sciences 103, 8066–8071 (2006).

10. S. Khan, M. Dapice, T. S. Reese, Effects of mot gene expression on the structure of the flagellar motor. Journal of Molecular Biology 202, 575–584 (1988).

11. A. Paulick et al., Two different stator systems drive a single polar flagellum in *Shewanella oneidensis* MR-1. Mol Microbiol 71, 836–850 (2009).

12. M. Ito et al., MotPS is the stator-force generator for motility of alkaliphilic *Bacillus*, and its homologue is a second functional Mot in *Bacillus subtilis*. Mol Microbiol 53, 1035–1049 (2004).

13. M. Schniederberend et al., Modulation of flagellar rotation in surface-attached bacteria: A pathway for rapid surface-sensing after flagellar attachment. PLOS Pathogens 15, e1008149 (2019).

14. A. L. Hook et al., Simultaneous tracking of *Pseudomonas aeruginosa* motility in liquid and at the solid-liquid interface reveals differential roles for the flagellar stators. mSystems 4, e00390–00319 (2019).

15. A. E. Baker et al., Flagellar Stators Stimulate c-di-GMP Production by *Pseudomonas aeruginosa*. Journal of Bacteriology 201, e00741–00718 (2019).

16. C. M. Toutain, M. E. Zegans, G. A. O’Toole, Evidence for two flagellar stators and their role in the motility of *Pseudomonas aeruginosa*. Journal of bacteriology 187, 771–777 (2005).

17. F. Wang, J. Yuan, H. C. Berg, Switching dynamics of the bacterial flagellar motor near zero load. Proceedings of the National Academy of Sciences 111, 15752–15755 (2014).

18. P. P. Lele, B. G. Hosu, H. C. Berg, Dynamics of mechanosensing in the bacterial flagellar motor. Proceedings of the National Academy of Sciences 110, 11839–11844 (2013).

19. A. Zöttl, J. M. Yeomans, Enhanced bacterial swimming speeds in macromolecular polymer solutions. Nature Physics 15, 554–558 (2019).

20. S. L. Kuchma et al., Cyclic di-GMP-mediated repression of swarming motility by *Pseudomonas aeruginosa* PA14 requires the MotAB stator. Journal of Bacteriology 197, 420–430 (2015).

21. C. M. Toutain, N. C. Caizza, M. E. Zegans, G. A. O’Toole, Roles for flagellar stators in biofilm formation by *Pseudomonas aeruginosa*. Research in Microbiology 158, 471–477 (2007).

22. Jacinta C. Conrad et al., Flagella and pili-mediated near-surface single-cell motility mechanisms in *P. aeruginosa*. Biophysical Journal 100, 1608–1616 (2011).

23. A. S. Utada et al., *Vibrio cholerae* use pili and flagella synergistically to effect motility switching and conditional surface attachment. Nature Communications 5, 4913 (2014).

24. J. de Anda et al., High-speed “4D” computational microscopy of bacterial surface motility. ACS Nano 11, 9340–9351 (2017).

25. Y. Yuan, K. VanderWerf, M. D. Shattuck, C. S. O’Hern, Jammed packings of 3D superellipsoids with tunable packing fraction, coordination number, and ordering. Soft Matter 15, 9751–9761 (2019).

26. A. Boromand, A. Signoriello, F. Ye, C. S. O’Hern, M. D. Shattuck, Jamming of Deformable Polygons. Physical Review Letters 121, 248003 (2018).

27. K. VanderWerf, A. Boromand, M. D. Shattuck, C. S. O’Hern, Pressure dependent shear response of jammed packings of frictionless spherical particles. Physical Review Letters 124, 038004 (2020).

28. Q. Wu, T. Bertrand, M. D. Shattuck, C. S. O’Hern, Response of jammed packings to thermal fluctuations. Physical Review E 96, 062902 (2017).

29. D. van Ditmarsch et al., Convergent evolution of hyperswarming leads to impaired biofilm formation in pathogenic bacteria. Cell Reports 4, 697–708 (2013).

30. G. Natan, V. M. Worlitzer, G. Ariel, A. Be’er, Mixed-species bacterial swarms show an interplay of mixing and segregation across scales. Scientific Reports 12, 16500 (2022).

31. K. A. Lewis et al., Nonmotile subpopulations of *Pseudomonas aeruginosa* repress flagellar motility in motile cells through a type IV pilus- and Pel-dependent mechanism. Journal of Bacteriology 204, e00528–00521 (2022).

32. K. VanderWerf, W. Jin, M. D. Shattuck, C. S. O’Hern, Hypostatic jammed packings of frictionless nonspherical particles. Phys Rev E 97, 012909 (2018).

33. Y.-G. Tao, W. K. d. Otter, J. T., Padding, J. K. G., Dhont, W. J. Briels, Brownian dynamics simulations of the self- and collective rotational diffusion coefficients of rigid long thin rods. The Journal of Chemical Physics 122, 244903 (2005).

34. E. R. Weeks, J. C. Crocker, A. C. Levitt, A. Schofield, D. A. Weitz, Three-dimensional direct imaging of structural relaxation near the colloidal glass transition. Science 287, 627–631 (2000).

35. C. S. O’Hern, L. E. Silbert, A. J. Liu, S. R. Nagel, Jamming at zero temperature and zero applied stress: The epitome of disorder. Physical Review E 68, 011306 (2003).

36. K. E. Boyle et al., Metabolism and the evolution of social behavior. Molecular Biology and Evolution 34, 2367–2379 (2017).

37. J. Yan, H. Monaco, J. B. Xavier, The Ultimate Guide to Bacterial Swarming: An Experimental Model to Study the Evolution of Cooperative Behavior. Annual Review of Microbiology 73, 293–312 (2019).

38. M. Deforet, D. van Ditmarsch, C. Carmona-Fontaine, J. B. Xavier, Hyperswarming adaptations in a bacterium improve collective motility without enhancing single cell motility. Soft Matter 10, 2405–2413 (2014).

39. M. E. Anyan et al., Type IV pili interactions promote intercellular association and moderate swarming of *Pseudomonas aeruginosa*. Proceedings of the National Academy of Sciences 111, 18013–18018 (2014).

40. I. Grobas, M. Polin, M. Asally, Swarming bacteria undergo localized dynamic phase transition to form stress-induced biofilms. eLife 10, e62632 (2021).

41. C. B. Whitchurch, T. Tolker-Nielsen, P. C. Ragas, J. S. Mattick, Extracellular DNA required for bacterial biofilm formation. Science 295, 1487–1487 (2002).

42. Z. Xing et al., Microrheology of DNA hydrogels. Proceedings of the National Academy of Sciences 115, 8137–8142 (2018).

43. S. C. Chew et al., Dynamic remodeling of microbial biofilms by functionally distinct exopolysaccharides. mBio 5, e01536–01514 (2014).

44. W. B. Russel, N. J. Wagner, J. Mewis, Divergence in the low shear viscosity for Brownian hard-sphere dispersions: At random close packing or the glass transition? Journal of Rheology 57, 1555–1567 (2013).

45. B. R. Parry et al., The bacterial cytoplasm has glass-like properties and is fluidized by metabolic activity. Cell 156, 183–194 (2014).

46. A. E. Baker et al., PilZ domain protein FlgZ mediates cyclic di-GMP-dependent swarming motility control in *Pseudomonas aeruginosa*. J Bacteriol 198, 1837–1846 (2016).

47. S. L. Kuchma et al., BifA, a cyclic-Di-GMP phosphodiesterase, inversely regulates biofilm formation and swarming motility by *Pseudomonas aeruginosa* PA14. J Bacteriol 189, 8165–8178 (2007).

48. J. J. Harrison et al., Elevated exopolysaccharide levels in *Pseudomonas aeruginosa* flagellar mutants have implications for biofilm growth and chronic infections. PLOS Genetics 16, e1008848 (2020).

49. L. Hou, A. Debru, Q. Chen, Q. Bao, K. Li, AmrZ regulates swarming motility through cyclic di-GMP-dependent motility inhibition and controlling Pel polysaccharide production in *Pseudomonas aeruginosa* PA14. Frontiers in Microbiology 10 (2019).

50. L. McCarter, M. Hilmen, M. Silverman, Flagellar dynamometer controls swarmer cell differentiation of *V. parahaemolyticus*. Cell 54, 345–351 (1988).

51. T. Julou et al., Cell-cell contacts confine public goods diffusion inside *Pseudomonas aeruginosa* clonal microcolonies. Proceedings of the National Academy of Sciences 110, 12577–12582 (2013).

52. C. R. Armbruster et al., Correction: Heterogeneity in surface sensing suggests a division of labor in *Pseudomonas aeruginosa* populations. eLife 9, e59154 (2020).

53. C. K. Lee et al., Social cooperativity of bacteria during reversible surface attachment in young biofilms: a quantitative comparison of *Pseudomonas aeruginosa* PA14 and PAO1. mBio 11, e02644–02619 (2020).

54. C. K. Lee et al., Multigenerational memory and adaptive adhesion in early bacterial biofilm communities. Proceedings of the National Academy of Sciences 115, 4471–4476 (2018).

55. H. P. Schweizer, Allelic exchange in *Pseudomonas aeruginosa* using novel ColE1-type vectors and a family of cassettes containing a portable oriT and the counter-selectable *Bacillus subtilis* sacB marker. Mol Microbiol 6, 1195–1204 (1992).

56. S. L. Kuchma, E. F. Griffin, G. A. O’Toole, Minor pilins of the type IV Pilus system participate in the negative regulation of swarming motility. Journal of Bacteriology 194, 5388–5403 (2012).

57. D. G. Ha, S. L. Kuchma, G. A. O’Toole, Plate-based assay for swarming motility in *Pseudomonas aeruginosa*. Methods Mol Biol 1149, 67–72 (2014).

58. D. G. Ha, S. L. Kuchma, G. A. O’Toole, Plate-based assay for swimming motility in *Pseudomonas aeruginosa*. Methods Mol Biol 1149, 59–65 (2014).

